# Argonomics Explorer: an interactive database for dissecting Argonaute-RNA regulatory networks

**DOI:** 10.64898/2026.11.25.769154

**Authors:** Wei-Sheng Wu, Mira Bhandari, Chun-Yuan Tai, Shou-Wei Huang, Heng-Chi Lee

**Affiliations:** Department of Electrical Engineering, National Cheng Kung University, Tainan 701, Taiwan; Department of Molecular Genetics and Cell Biology, University of Chicago, Chicago, IL, 60637, USA

**Keywords:** Argonaute, small RNA, miRNA, piRNA, gene regulation

## Abstract

Argonaute (AGO) family proteins associate with small RNAs (sRNAs) to form complexes that direct sequence-specific regulation of target RNAs. Recent studies in *Caenorhabditis elegans, Drosophila melanogaster,* and human cell lines have systematically characterized the repertoire of sRNA associations and targets for individual AGOs in these organisms; however, the lack of an accessible, searchable platform limits efficient comparison of the specificity and redundancy of Argonaute functions. Here, we present Argonomics Explorer (AE), an interactive database suite that integrates publicly available sRNA sequencing datasets to enable systematic interrogation of AGO-sRNA and AGO-target interactions across common model systems. AE combines independent AGO-IP datasets, AGO mutant-derived sRNA profiles, and systematic analyses to provide a cross-validated framework for examining AGO-RNA interactions. This database empowers users to query specific AGOs, transcripts, or sRNAs, and to compare associated sRNAs or targets across AGOs. Specifically, we recapitulate findings that certain microRNA (miRNA) star strands are uniquely loaded onto AGO-2 but not AGO-1 in flies. In addition, we report a preferential association of most members of the mir-35 family with the worm Argonaute ALG-2 and the association of the oncogenic oncomiR-1 miRNA superfamily with human AGO2. Such applications illustrate how the AE framework facilitates investigations into AGO specificity, redundancy, and the broader regulatory networks that shape gene expression. The Argonomics Explorer suite is freely available at https://cosbi10.ee.ncku.edu.tw/AE/.

## Introduction

Argonaute proteins (AGOs) are central effectors of small RNA-guided gene regulatory mechanisms that are conserved among nearly all eukaryotes (Seroussi et al. 2023; Fusca et al. 2025). Upon loading of a small RNA (sRNA) guide, an AGO forms an RNA-induced silencing complex (RISC) that can associate with corresponding target RNAs through complementary base pairing. This interaction enables the RISC to regulate target gene expression through various mechanisms such as repressing translation of mRNAs, promoting their degradation or directing transcriptional silencing (Hutvagner and Simard 2008).

Despite sharing common structural features, diverse AGOs operate in specialized sRNA-guided regulatory pathways, many of which are broadly conserved in metazoans. For instance, the small interfering RNA (siRNA) and PIWI interacting RNA (piRNA) pathways act as gene silencing mechanisms against foreign or selfish RNAs from viruses or transposons (Jouravleva and Zamore 2025). The microRNA (miRNA) pathway also plays a widely conserved role in fine-tuning the expression of endogenous RNAs. In *C. elegans,* RDE-1 is the effector of RNAi, while AGOs including ALG-1, ALG-2, and ALG-5 are key effectors of miRNA-mediated gene silencing (Yigit et al. 2006; Pal et al. 2024). The two *Drosophila* AGOs are similarly partitioned by pathway with AGO2 functioning in the siRNA pathway and AGO1 acting as the primary effector of the miRNA pathway (Meyer et al. 2006). The human AGO family consists of four paralogs, AGO1-4, which are broadly co-expressed and thought to be largely redundant regulators of the miRNA pathway (Nakanishi 2022). Nonetheless, multiple exceptions to these generally accepted AGO-sRNA pathway associations have been reported. For example, some worm miRNAs are associated with the siRNA AGO RDE-1 (Seroussi et al. 2023), and multiple fly miRNA* strand species are sorted to the siRNA AGO, AGO2 (Okamura et al. 2009). In addition, the worm genome exhibits extensive expansion of AGO family members that associate with endogenous siRNAs and amplified sRNAs that are enriched in various cell types (Seroussi et al. 2023; Liu et al. 2023; Fusca et al. 2025). Thus, the specificity and redundancy of Argonaute proteins remain an area of active investigation across organisms.

Two recent studies systematically characterized the sRNA network of 19 *C. elegans* AGOs by generating small RNA sequencing libraries using two orthologous approaches: immunoprecipitation of 3X-FLAG-tagged AGO complexes and sRNA extraction from systematically deleted AGO mutants (Seroussi et al. 2023; Liu et al. 2023). Analogous immunoprecipitation studies have also been conducted by Kingston and Bartel (2021) in *Drosophila melanogaster* for FLAG-tagged Argonaute clade members, AGO1 and AGO2 and in human cells for AGO-1 to AGO-4 (Figure 1A; Dueck et al. 2012). Notably, these datasets from flies, humans, and worms include input sRNA sequencing data, allowing for the determination of enriched AGO binding against baseline cellular abundance. Collectively, while these works provide valuable data characterizing the specificity and redundancy of AGOs, efficient exploration of these datasets is currently limited by the absence of a searchable and straightforward platform to navigate through AGO-RNA interactions of interest (Dueck et al. 2012; Kingston and Bartel 2021; Seroussi et al. 2023; Liu et al. 2023).

**Figure 1.**
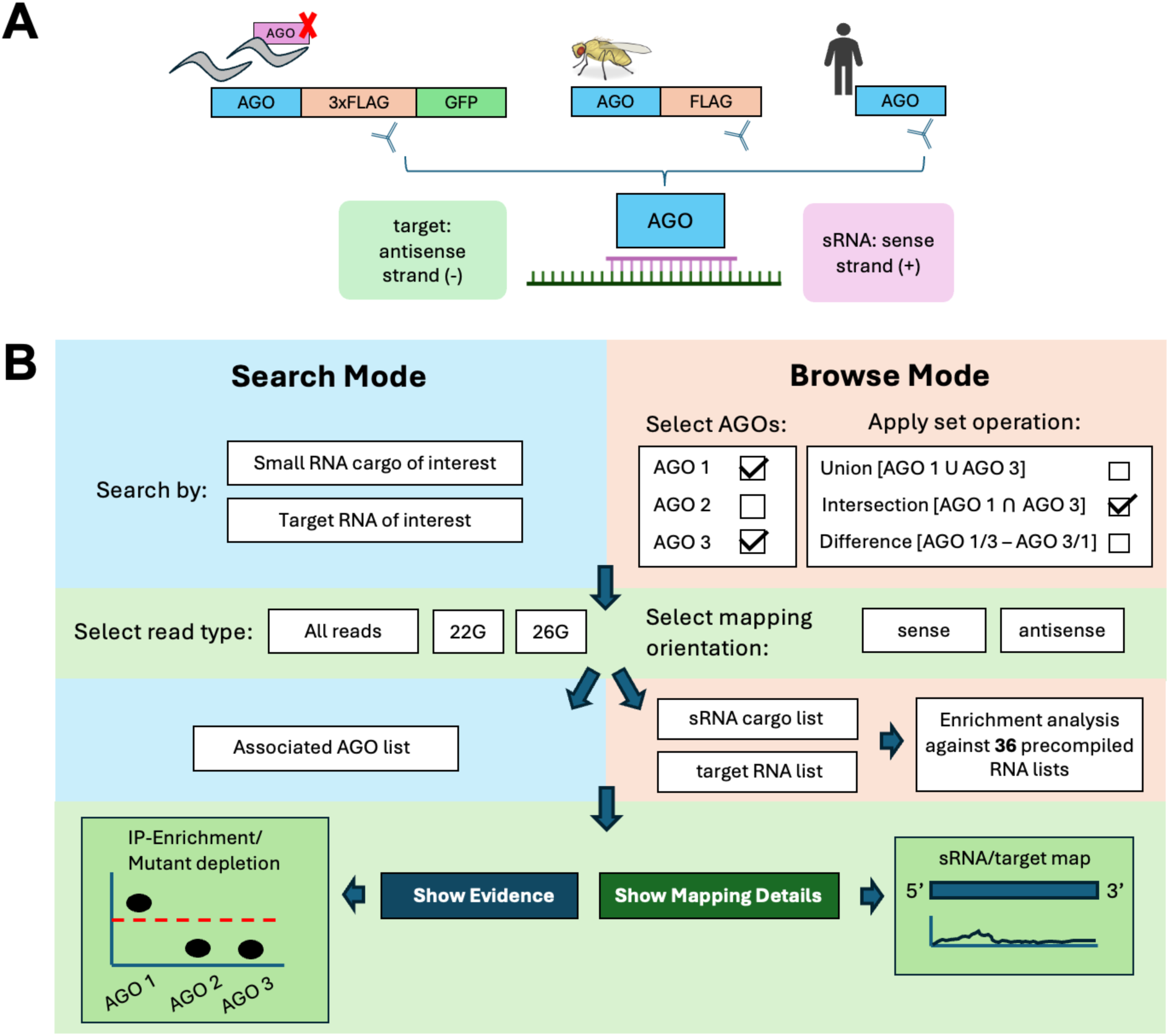
Argonomics Explorer workflow. sRNA-sequencing library acquisition and mapping strategies for *C. elegans, Drosophila,* and human AGOs (**1A**). User guide for navigating AE databases in “Search” or “Browse” modes (**1B**).

To this end, we have developed the Argonomics Explorer suite, an interactive collection of databases comprised of WAE, HAE, and FAE for *C. elegans, Drosophila,* and human Argonautes, respectively. This suite enables systematic exploration of Argonaute regulatory networks and rapid identification and comparison of AGO-associated small RNAs and AGO-RNA interactions across datasets (Figure 1B). The sense mapping of these data identifies AGO-associated or AGO-dependent small RNAs, whereas antisense mapping identifies potential target mRNAs. AE focuses on target RNAs that exhibit extensive sequence complementarity (fewer than two mismatches) to their associated sRNAs. Targets mediated by limited or partial complementarity, such as many canonical miRNA targets, are thus beyond the primary scope of this database. Users are therefore encouraged to consult specialized resources or databases for exploring these interactions (Vlachos et al. 2014; Lewis et al. 2003; Yang et al. 2025).

The database provides two querying pipelines: first, users may input a particular sRNA or RNA target of interest and retrieve a catalog of IP-enriched and/or mutant-depleted AGO interactors. Alternatively, multiple AGOs can be selected to generate and compare lists of sRNAs or complementary RNA targets, and to identify unique or shared interactor lists among AGOs. Additionally, users are provided with details including fold-change data from the original IP experiments, read mapping distributions illustrating AGO binding footprints, and the ability to cross-reference sRNAs and targets spanning multiple datasets. This approach also remains agnostic to prior annotations by mapping all sRNA-seq reads to the complete reference transcriptome. Below, we present several examples highlighting the usage of AE. Taken together, the Argonomics Explorer database suite transforms expansive datasets by delivering an accessible resource for supporting research on AGO-mediated interactions across model systems, enabling high-level browsing and acting as a hypothesis generator.

## Results

Each organism-specific database within the AE suite supports two querying modes that enable users to extract either the AGOs that associate with an sRNA or target of interest (Search mode) or, conversely, the target genes that are regulated by or sRNAs associated with AGO(s) of interest (Browse mode). The first mode can be accessed from the Search page (Figures 2-3). This page allows users to search by gene name or transcript ID and view the AGOs that associate with the RNA of interest. As reads mapping in both sense and anti-sense directions are retained in AE, users can either search for a specific sRNA such as a miRNA (e.g. *lin-4* miRNA) or piRNA (e.g. 21ur-4863) to retrieve AGOs enriched or mutant-depleted for corresponding reads in the sense direction (Figure 2). Alternatively, users can query a target transcript of interest (e.g. piRNA targets *xol-1* or *fbxb-97* mRNA) which will instead retrieve associated AGOs based on reads mapped in the antisense direction (Figure 3).

**Figure 2.**
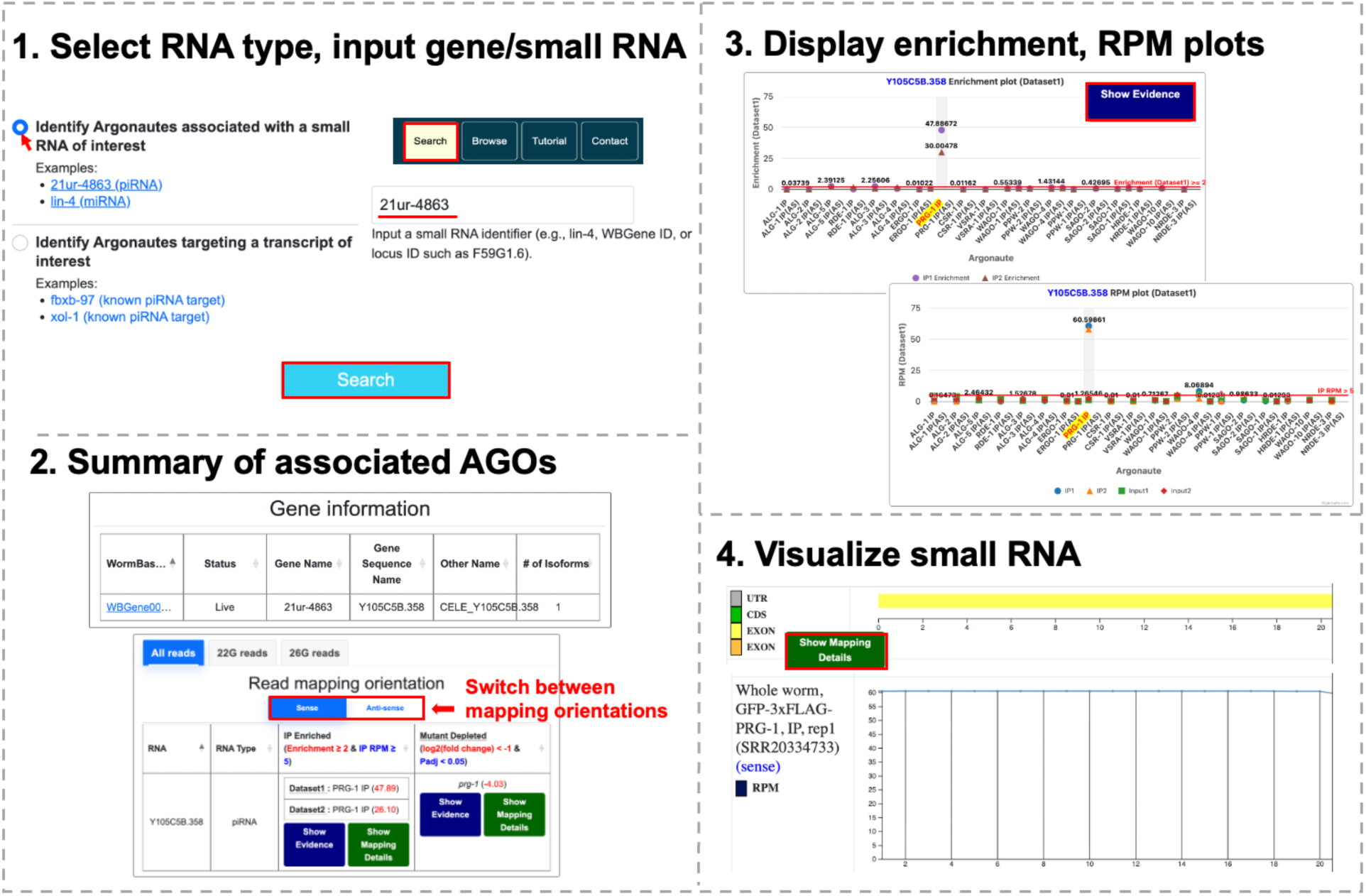
Walkthrough of search-by-sRNA querying pipeline using piRNA, 21ur-4863. The “Search” page allows users to query AGO associations for either sRNA cargo (sense reads) or a regulatory target (antisense reads) (**2.1**). The results page displays basic cataloguing information corresponding to the sRNA input in addition to a summary of the AGO-IPs in which association with the queried sRNA is significantly enriched/depleted (**2.2**). Clicking “Show Evidence” reveals detailed statistical plots showing IP-enrichment and RPM values for each AGO. (**2.3**) Selecting “Show Mapping Details” displays a graphical view of the selected sRNA, showing the positional distribution of mapped reads (**2.4**).

**Figure 3.**
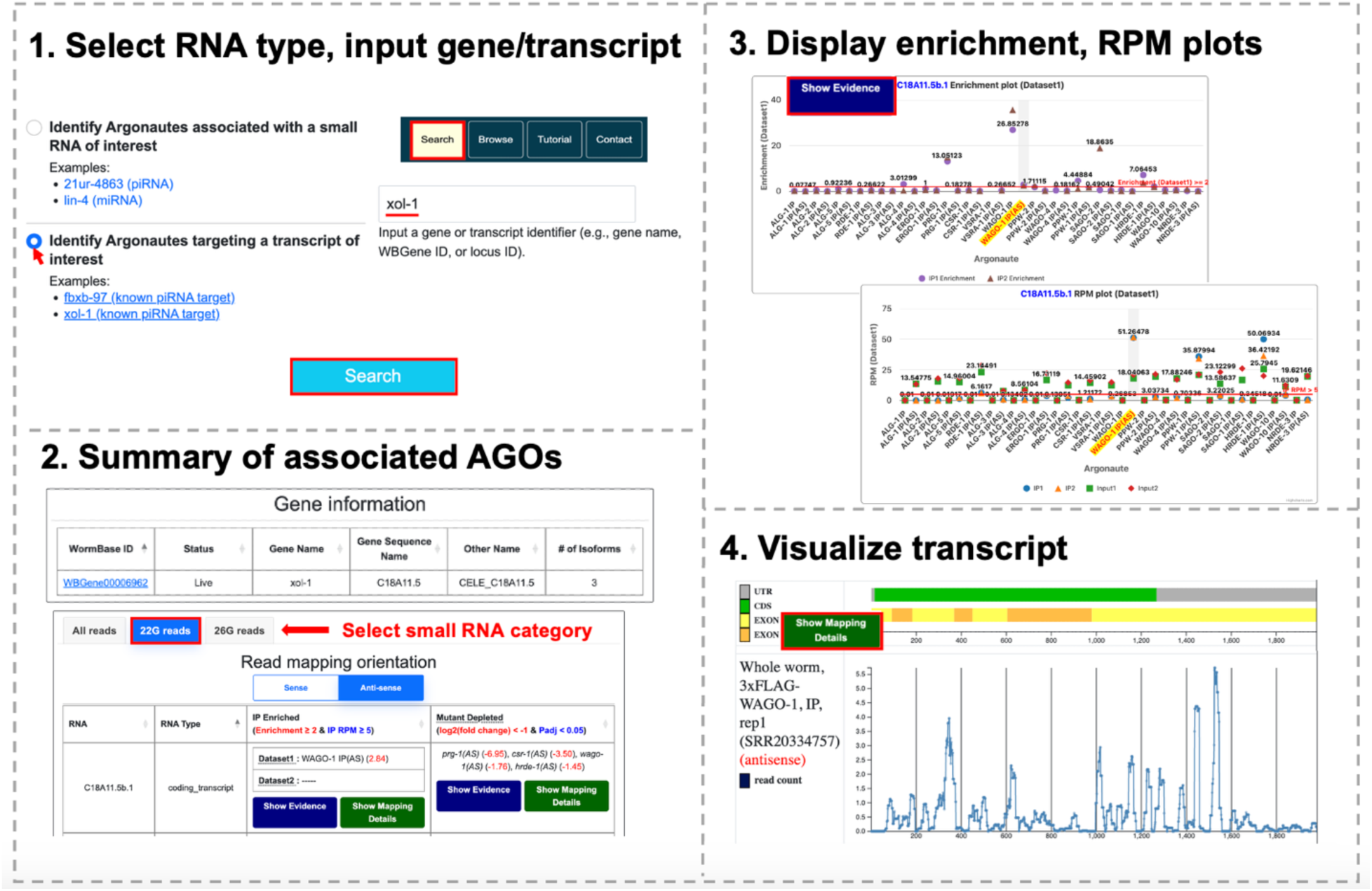
Walkthrough of search-by-target querying pipeline using *xol-1.* Here, users can search for a gene target via name, WBGene ID, or locus ID (**3.1**). The two resulting tables describe gene aliases and isoforms in addition to a summary of the AGO-IPs in which the target is enriched in both datasets which can be further filtered by type of sRNA (**3.2**). Clicking “Show Evidence” reveals detailed statistical plots showing IP-enrichment and RPM values for each AGO. (**3.3**) Selecting “Show Mapping Details” displays a graphical view of the selected transcript, showing its structural features (UTR, CDS, exons) and the positional distribution of mapped datasets (**3.4**).

The second querying mode available through AE can be accessed from the Browse page and enables users to retrieve lists of sRNAs or targets associated with AGO(s) of interest (Figure 4-5). Similarly to search mode, here, users can either retrieve lists of AGO-associated sRNA from sense reads or target transcripts derived from antisense reads. Users can select and compare multiple Argonautes simultaneously to identify sRNAs or target RNAs that are uniquely or commonly associated with different Argonautes, thereby revealing patterns of Argonaute specificity and functional redundancy. The resulting list and associated data can be filtered based on RNA type (such as mRNA, non-coding RNA, transposon mRNA, etc.) and exported directly from AE. In *C. elegans* Browse mode, users can also access an over/under-representation analysis table that summarizes whether the RNAs retrieved from a query are enriched or depleted for RNAs belonging to 36 precompiled transcript type datasets including piRNAs, miRNAs, or coding transcripts (Table S1). This table is automatically generated when a query is submitted in the Browse page.

**Figure 4.**
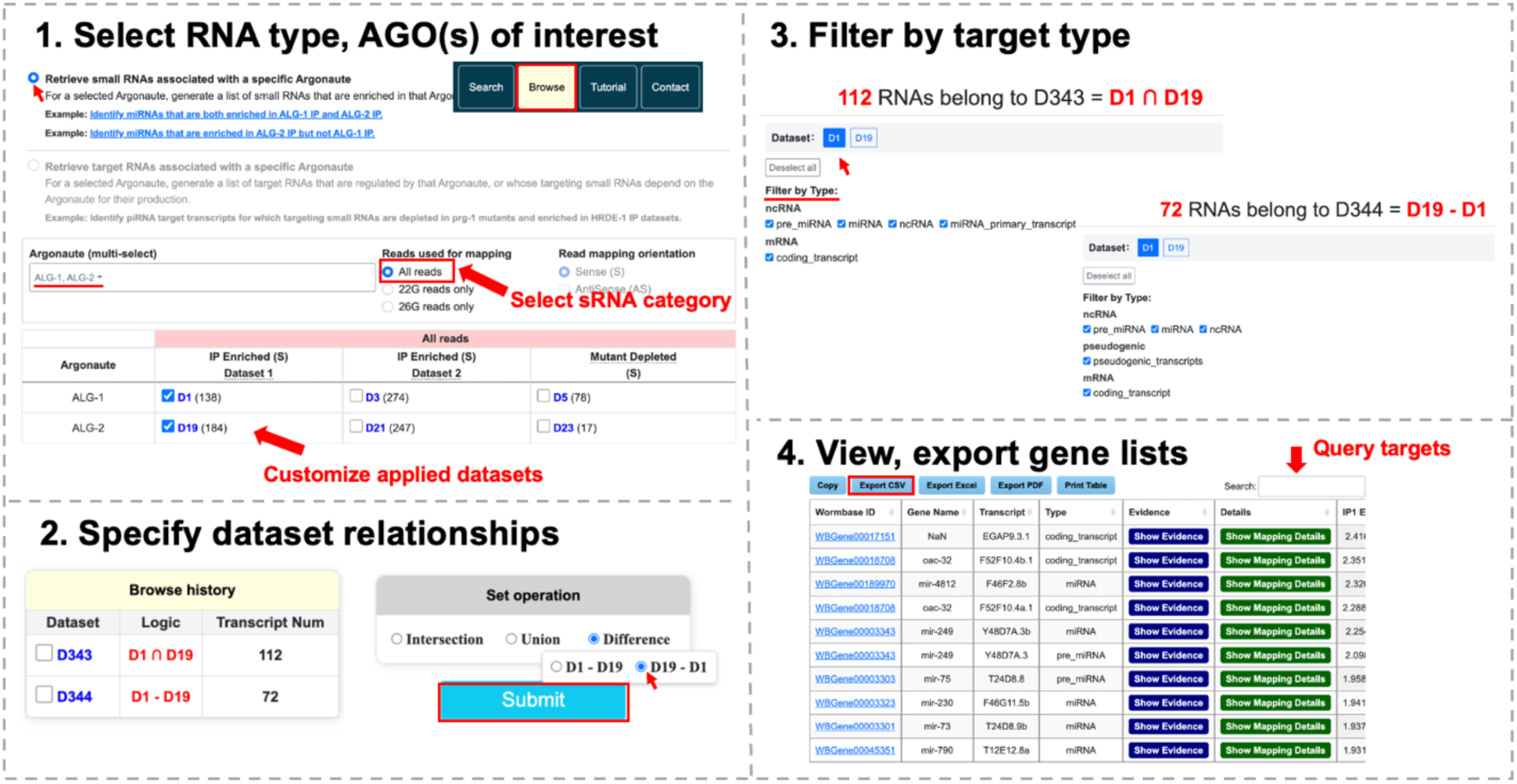
Generating comparative ALG-1, ALG-2 target lists in Browse mode. Specify AGOs of interest, datasets considered and reads used for mapping (**4.1**). Apply set operations to selected datasets, retaining access to past queries (**4.2**). Select transcript categories of interest and choose display dataset for resulting table (**4.3**). View and export target tables containing IP enrichment, IP and input abundance (RPM) from specified dataset (**4.4**).

**Figure 5.**
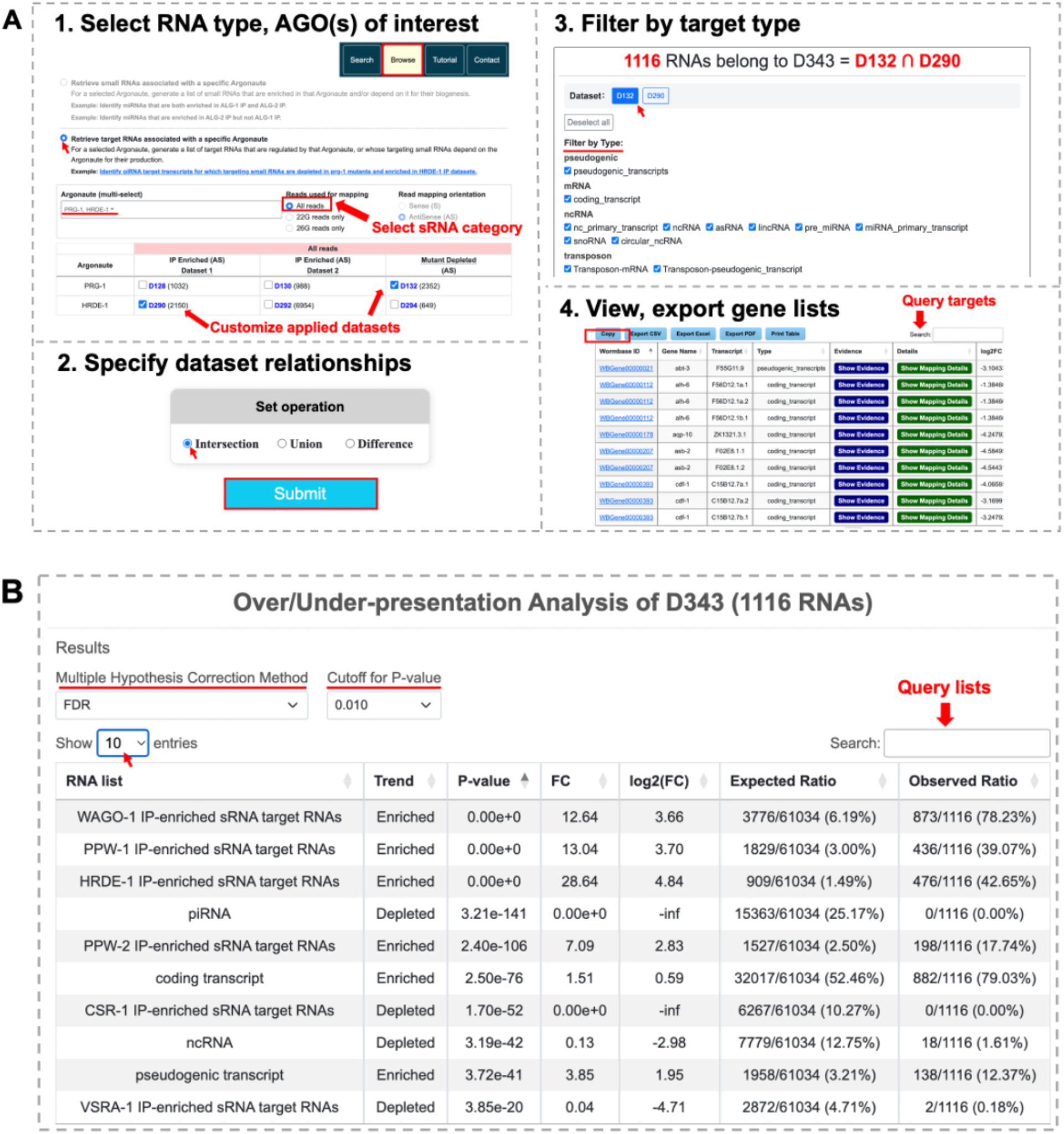
Example 2 walkthrough using the “Browse” page to generate target lists enriched in HRDE-1 IP and depleted in *prg-1* mutants. Users can choose to generate lists of sRNA cargo or regulatory targets based on sRNA classification for single or multiple AGOs. These selections generate a pool of relevant datasets that can be filtered further by the user to specify consideration of one or both datasets, or mutant depleted vs. IP datasets (**5A.1**). The resulting gene list for multiple AGOs can also be customized by applying a set operation (**5A.2**). The results page displays the number of RNA targets belonging to the queried dataset which can in turn be filtered by target RNA type (**5A.3**). When all user customizations have been made, the final target list including enrichment/depletion values and basic gene information for each target can be copied, printed, or exported in.csv, .xlsx, or .pdf forms (**5A.4**). Example over/under-representation analysis table generated automatically upon making a query in WAE’s Browse mode (**5B**).

The following examples illustrate detailed uses of both Search and Browse modes and demonstrate how the AE database can be harnessed to generate biologically meaningful hypotheses about AGO-RNA regulatory relationships across common model systems.

### Search Mode Example 1: Identify *C. elegans* AGOs associated with the piRNA, 21ur-4863

The piRNA 21ur-4863 is reported to cooperatively silence expression of the developmental regulator, *xol-1,* mRNA. Deletion of 21ur-4863 yields upregulation of *xol-1* mRNA and protein, emphasizing this piRNA’s functional relevance to dosage compensation and sex determination (Tang et al. 2018; Shen et al 2018). This well-defined regulatory role makes 21ur-4863 a useful case for asking which AGOs it associates with.

To query 21ur-4863, the user can navigate to the “Search” page and select the search option specific to sRNAs, i.e. reads mapped in the sense orientation. After typing 21ur-4863 into the prompt box and clicking search, two sets of information are presented (Figure 2, Box 1). First, a “Gene Information” table provides a core report for the matched gene, including its WormBase ID, name, aliases, and number of associated isoforms (slice variants, precursors). The second table summarizes IP-enrichment and mutant-depletion data for each isoform of the queried sRNA or target (Figure 2, Box 2). As expected, by searching for AGOs that associate with 21ur-4863, WAE reveals enrichment of its reads in the PIWI AGO, PRG-1 IP and depletion in *prg-1* mutants, consistent with the established role of PRG-1 as the primary AGO responsible for piRNA loading in *C. elegans* (Batista et al. 2008).

Users can further access the underlying enrichment/depletion values across AGOs by selecting the “Show Evidence” option, opening a new tab displaying RPM and fold change plots of these values for each AGO and allowing users to view data corresponding to AGOs beyond the threshold limits specified in the summary table (Figure 2, Box 3). Furthermore, separate plots are provided for IP enrichment corresponding to Dataset 1 and 2, enabling comparison between the two independent data sources. Clicking “Show Mapping Details” allows users to visualize the distribution of mapped reads along the queried RNA (Figure 2, Box 4).

### Search Mode Example 2: Search for *C. elegans* AGOs that target the sex-determining factor, *xol-1*

In *C. elegans,* piRNAs trigger silencing by initiating RdRP-dependent synthesis of secondary sRNAs that are antisense to the target mRNA. These secondary sRNAs, known as 22G-RNAs for their length and first nucleotide preference, are loaded onto WAGO-clade AGOs, including WAGO-1 and HRDE-1/WAGO-9 to silence piRNA targets (Shirayama et al. 2012; Lee et al. 2012). As previously described, the master sex-determination switch, *xol-1,* is a well-characterized piRNA target in *C. elegans*. Since its output is dose-sensitive, modest shifts in expression can reprogram developmental fate between hermaphrodite and male. Here, we use *xol-1* as an example to explore which AGOs associate with sRNAs that target *xol-1* mRNAs.

To query *xol-1* on the “Search” page, the user can select the second provided option to identify AGOs targeting a transcript of interest (Figure 3, Box 1). Making this selection will ensure that WAE returns enrichment/depletion for reads mapping in the antisense orientation by default. However, the user retains the ability to switch the mapping between sense and antisense after the query has been submitted. After entering *xol-1* into the prompt box, the results are returned in the same two-table format described in Search Mode Example 1. The “Gene Information” table provides annotation details for the matched gene while the accompanying summary table reports IP-enrichment and mutant-depletion data for *xol-1* across all profiled AGOs surpassing the required thresholds for enrichment or depletion (Figure 3, Box 2). Implementing this pipeline on *xol-1,* WAE reveals that *xol-1* reads are enriched in WAGO-1 and HRDE-1IPs and are depleted in *prg-1, csr-1, wago-1,* and *hrde-1* mutants. Selecting “Show Evidence” opens a detailed view of RPM and fold-change values, enabling users to assess the strength of AGO associations and inspect data from both datasets independently (Figure 3, Box 3). “Show Mapping Details” can then be selected to visualize the positional context of the detected AGO associations across the *xol-1* locus, guiding the identification of potential AGO targeting hotspots.

This output supports the notion of a multi-step regulatory cascade for *xol-1* and is corroborated by previous findings that disruption of the WAGO-mediated RNAi pathway leads to aberrantly enhanced *xol-1* mRNA expression and that *xol-1* is regulated by PRG-1 and its associated piRNAs including 21ur-4863 and 21ux-1 (Tang et al. 2018; Shen et al 2018). According to WAE, *xol-1* reads are also significantly depleted in *csr-1* mutant backgrounds, providing a basis for future exploration of a role for CSR-1, an AGO uniquely known to promote expression or fine-tuning of its targets, as contributing to the proper biogenesis or stabilization of *xol-1-*targeting sRNAs (Cecere et al, 2014; Charlesworth et al, 2021).

### Browse Mode Example 1: Generate comparative lists of miRNAs associated with *C. elegans* ALG-1 and ALG-2

The ability of WAE to generate comprehensive lists of targets that are enriched in particular AGO pulldowns or depleted when an AGO’s function is lost enables dissection of the regulatory roles of key AGOs across various sRNA pathways. This is exemplified by the “Browse” querying pipeline, which we use here to generate comparative lists of miRNAs that are associated with ALG-1 and ALG-2.

Although two *C. elegans* miRNA AGOs, ALG-1 and ALG-2, are paralogs that share over 75% of their amino acid makeup and similar spatiotemporal expression patterns during development, recent evidence suggests they are not fully redundant (Aalto et al. 2018). For example, single mutants of either *alg-1* or *alg-2* yield defects in brood size and number of oocytes, implying that one functional miRNA AGO-encoding gene is not sufficient to compensate for the loss of the other (Brown and Montgomery 2018).

To perform these queries, the user can navigate to the “Browse” page and select the option to retrieve sRNAs associated with a specific Argonaute. The generated gene list can consist of sRNAs for a specific AGO or combination of AGOs, which can be specified using the multi-select feature in the AGO prompt box. (Figure 4, Box 1). Like the search-by-transcript querying pipeline, the category of sRNA reads (which defaults to all reads) considered for the gene list can also be specified here. These selections will populate a table with the datasets relevant to each selected AGO that can be filtered further by the user. The datasets corresponding to ALG-1 and ALG-2 IP enrichment, D1 and D19, which contain 138 and 184 sRNAs, respectively. To identify sRNAs associated with both ALG-1 and ALG-2, the datasets D1 and D19 can be selected from the table, and the intersection set can be selected to retain transcripts enriched in both AGO pulldowns (Figure 4, Box 2). Submitting this query generates an RNA list representing the shared 112 sRNAs, among which many are miRNAs. To identify transcripts preferentially associated with ALG-2, the user can return to the dataset selection table, retain the same AGO and dataset selections, and switch the set operation to difference. Selecting D19 – D1 will specify sRNAs enriched in ALG-2 IP but not in ALG-1 IP. The resulting dataset consists of 72 sRNAs. The third available set operation in WAE is the union option, which will retain all transcripts that are present in either one of these datasets or both of them (Figure 4, Box 2).

The results first specify the number of RNAs belonging to the retained data given the data pools and set operation chosen by the user (Figure 4, Box 3). These RNAs can be further filtered based on categories like pseudogenic transcripts, miRNA, ncRNA, or transposons, which will consequently update the target RNA count shown below the table. The resulting gene table will include core information for each target including gene name and RNA type as well as enrichment, depletion, or normalized read count values for each replicate performed in the original studies (Figure 4, Box 4). Any of these metrics can be used to sort the list by clicking on the column heading once for an ascending order sort and twice for descending order. Additionally, individual targets of interest can be queried via a search box above the target table. Once the table has been filtered and sorted based on user needs, it can be exported in publication-ready formats or plain text format (.csv) for use in downstream bioinformatic analyses (Table S2).

Querying for targets enriched in ALG-2 IP but not ALG-1 IP returns 72 RNAs without transcript type filters and 34 miRNAs when the mature miRNA type is specified. Among these ALG-2 associated miRNAs, 7 out of 8 members of the *mir-35-42* miRNA cluster sharing the same seed region are represented (Supplementary Figure S1). Notably, *mir-35-41,* the miRNAs from this cluster that associate preferentially with ALG-2, are co-transcribed from a single primary transcript. On the other hand, *mir-42,* which is transcribed from a different locus, is associated with both ALG-1 and ALG-2. With these findings, WAE provides support for the hypothesis that genomic locus or primary transcript context may play a role in the sorting of worm miRNAs.

### Browse Mode Example 2: Identify transcripts depleted in *C. elegans* PIWI mutants and enriched in HRDE-1 IP

The PIWI Argonaute PRG-1, guided by piRNAs, performs broad genome surveillance and can trigger silencing of its targets (Lee et al. 2012). Once piRNAs identify their mRNA targets, the production of secondary sRNAs is triggered. These sRNAs are loaded onto WAGO-clade AGOs, specifically HRDE-1/WAGO-9, in the nucleus to enforce piRNA-mediated transcriptional silencing (Ding et al. 2023). Thus, we expect that the secondary sRNAs that recognize some piRNA targets will be depleted in *prg-1* mutants but enriched in HRDE-1 IP (Batista et al. 2008). Here, we use “Browse” mode in WAE to generate this piRNA target list.

To use WAE as a target list generator for this application, the user can again navigate to the “Browse” page of the database and select the second provided option to retrieve target RNAs associated with a specific Argonaute (Figure 5A, Box 1). PRG-1 and HRDE-1 can then be selected using the multi-select feature in the AGO prompt box. To identify *prg-1* mutant-depleted transcripts, the corresponding dataset, D132, is selected along with D290 to identify HRDE-1 IP-enriched targets derived from the same study, i.e. Dataset 1. The relationship between datasets can then be specified by selecting a set operation such as intersection, which generates a list of RNAs present in both the *prg-1* mutant-depleted and HRDE-1 IP-enriched datasets (Figure 5A, Box 2). After making these selections, the user can click submit to generate their gene list.

The *prg-*1-depleted, HRDE-1 IP-enriched table contains 1116 RNAs agnostic of RNA type and 882 targets when filtered for mRNA (Figure 5A, Box 3). Underneath the gene list table, users can find an option to generate over/under-representation tables for broad analysis of the transcript types present in the AGO IP-enriched target list (Figure 5A, Box 4). Users can select between False Discovery Rate (FDR) or Bonferroni correction and customize the P-value cutoffs for the enrichment/depletion analysis performed with the queried target list and precompiled transcript lists (Figure 5B). This analysis reveals that the transcripts belonging to the intersection of *prg-1* mutant-depleted and HRDE-1 IP-enriched reads also overlap significantly with WAGO-1 IP-enriched transcripts, which validates the canonical piRNA silencing cascade model where both WAGO-1 and HRDE-1 associate with secondary sRNAs to silence piRNA targets (Shirayama et al. 2012; Lee et al. 2012). The intersection operation can be applied iteratively to multiple sRNA or target lists, allowing users to identify RNAs that simultaneously satisfy multiple criteria and thereby progressively refine candidate sets based on increasingly specific regulatory features. In addition, it is also observed that these transcripts are enriched for coding transcripts and depleted for CSR-1 IP-enriched sRNA target RNAs. As CSR-1 is thought to antagonize the PRG-1/piRNA-mediated silencing pathway, it follows that these two pathways are operating on predominantly non-overlapping transcript sets (Charlesworth et al. 2021; Kirschner et al. 2025).

Together, these results demonstrate that WAE’s browse mode can rapidly generate coherent target lists that recapitulate established pathway logic, reveal genome-scale regulatory principles such as the mutual exclusivity of CSR-1 licensing and PRG-1/HRDE-1 silencing, and identify specific gene families or candidates for downstream analysis.

### Browse Mode Example 3: Assess preferential miRNA sorting to AGOs across species

A central question in AGO biology is whether AGOs predominantly associated with a single type of sRNA (siRNAs, piRNAs, or miRNAs) can load populations of other sRNA classes as well. This gap persists in different forms in each organism represented in the AE suite. In *C. elegans,* RDE-1 functions as the primary AGO in the exogenous RNAi pathway. This paradigm was challenged by the finding from Seroussi et al. (2023)’s IP-derived read data, which uncovered that RDE-1 binds specific miRNAs in addition to siRNAs. In *Drosophila,* the two strands of a miRNA precursor duplex are known to be asymmetrically sorted rather than co-loaded onto the same AGO (Okamura et al. 2009). The guide strand is preferentially retained by AGO1 to direct canonical miRNA-mediated silencing while the passenger miR* strand is instead selectively loaded onto AGO2, the primary effector of the siRNA pathway (Czech et al. 2008). This finding stands in contrast to previous models wherein miR* strands were presumed to be degraded byproducts of the miRNA maturation pathway (Kim et al. 2009).

Human AGO1-4 are thought to act with substantial functional redundancy in the miRNA pathway as all four paralogs are known to bind mature miRNAs to mediate translational repression (Landthaler et al. 2008). However, this redundancy model is complicated by the genetics of Argonaute syndrome, in which mutations in a single paralog, AGO2, were recently linked to Lessel-Krienkamp syndrome, a distinct neurodevelopmental disorder (Yang et al. 2025). This etiology suggests that at least some AGO2 targets may not be efficiently compensated by its three other paralogs.

Because AE Browse mode queries reads directly by AGO identity and RNA class without relying on prior pathway assignments, each of these examples can be recapitulated using the same underlying querying logic across WAE, FAE, and HAE modules.

Applied to *C. elegans,* querying against the IP reads of the siRNA AGO, RDE-1, from WAE Dataset 1 returns 172 miRNA-associated isoforms corresponding to 89 unique miRNAs out of 1111 total sRNAs (Figure 6). This indicates that a subset of endogenous miRNAs is indeed enriched in the RDE-1 pulldown despite RDE-1’s canonical designation as a dedicated siRNA Argonaute, recapitulating the findings from Seroussi et al. (2023) and creating a searchable list of the miRNA population associated with RDE-1.

**Figure 6.**
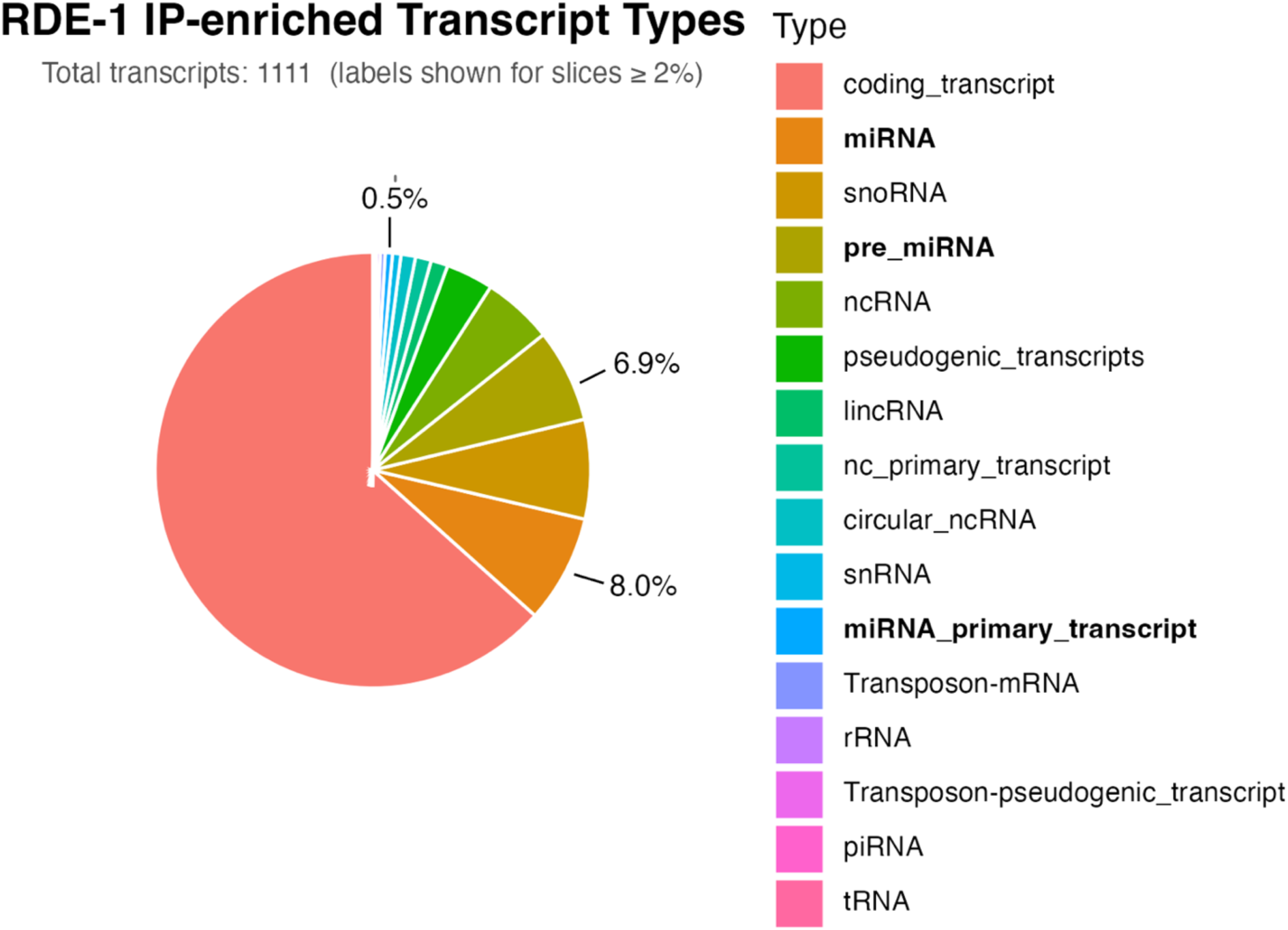
Transcript type composition of RDE-1 IP-enriched RNAs. Transcripts were classified by biotype using WormBase release, WS293. Percentages are shown for miRNA-associated biotypes (miRNA, pre_miRNA, and miRNA_primary_transcript; **bold** in legend).

In *Drosophila,* FAE additionally reports on an siRNA AGO, AGO2-IP dataset, in a *hen-1;dora* mutant background in which target-directed degradation (TDD) is blocked, providing a readout of steady-state AGO2 cargo that preserves lower-abundance or less stable species compared to wildtype AGO2. Querying the intersection of all three replicates of the AGO2-Hen1Dora IP dataset returns 25 miRNA-associated isoforms, of which every well-annotated locus corresponds to the miR* strand. When we apply the difference set operation (AGO2-Hen1Dora – AGO1-IP) to identify miR* species enriched in AGO2 but absent from the miRNA AGO AGO1-associated pool, FAE retrieves examples of selective loading that match previous reports, including bantam* and miR-184*, recapitulating the reported observations that selected miRNA* strands are sorted to AGO2 (Okamura et al. 2009 and Supplementary Figure S2).

In HAE, choosing AGO-1-4 as AGOs of interest in Browse mode and applying the difference operation to retain transcripts exclusively enriched in AGO-2-IPs returns a list of 385 RNAs. Filtering this list to the miRNA and mature miRNA categories and sorting by abundance, several of the most highly represented entries converge on the oncomiR-1 miRNA cluster and its paralogs, miR-17-92 and miR-106a-363 and miR-106b-25, which share the same seed region but originate from distinct primary miRNA (pri-miRNA) transcripts (Gruszka and Zakrzewska 2018). At the mature miRNA level, miR-92a-3p, miR-25-3p, and miR-106b-3p, originating from the oncomiR-1 pri-miRNA and the pri-miRNA transcripts for its two paralogs respectively, are enriched over two-fold compared to input. This pattern is absent from the other three AGO-IP datasets, indicating consistent, preferential AGO-2 loading across all three paralogous oncomiR-1 clusters (Supplementary Figure S3). Together, these examples demonstrate the utility of AE for identifying unique and shared AGO-associated miRNAs across organisms, providing a powerful hypothesis-generating framework for investigating AGO specificity, functional redundancy, and regulatory network architecture.

## Discussion

The Argonomics Explorer (AE) database suite was developed to address a growing gap between the scale of publicly available AGO-sRNA sequencing data and the tools available to analyze it in a systematic, cross-referenced manner. Beyond generating a compilation of these datasets, applying AE across three model systems to examine AGO-sRNA associations such as miRNA binding highlights that the boundaries drawn by pathway-based classifications of AGOs are more intertwined than the canonical framework of miRNA or siRNA-specific AGOs suggests. Specifically, AE recapitulates and produces lists of miRNAs that preferentially associate with the siRNA AGOs, RDE-1 in *C. elegans*, and *Drosophila* AGO2. WAE also identifies a distinct preference of the miR-35 family miRNAs for association with ALG-2 over ALG-1. Given that the miR-35 family plays an essential role in protecting *C. elegans* from radiation-induced apoptosis, the association of ALG-2 with the miR-35 family identified by AE provides a potential mechanistic explanation for the excess apoptosis observed in both *alg-2* and *mir-35-41* mutants following irradiation (Doll et al. 2019). Furthermore, AE reports unique enrichment of miRNA families with specific human AGOs, including the preferential association of AGO-2 with the oncomiR-1 superfamily of miRNAs found to be overexpressed in various lymphomas and solid tumors (Chakraborty and Krishnan 2017; Gruszka and Zakrzewska 2018). These results highlight that despite broad functional redundancy between human AGO-1-4, loading specificity exists at the level of individual miRNA families. Taken together, these examples underscore AE’s potential not only as a data retrieval and exploration tool, but also as a platform for probing the rules that govern AGO-cargo specificity and redundancy across organisms.

The utility of AE stems from three features that distinguish it from the static supplementary tables or single-purpose analysis scripts accompanying the original studies. First, by mapping all sRNA reads directly to reference transcriptomes rather than relying on the annotation conventions of the source publications, AE remains agnostic to prior classifications and uncovers noncanonical AGO associations through, for example, the identification of lists of miRNAs that associate with siRNA AGOs while maintaining the user’s ability to examine detailed mapping results manually. Second, AE provides a single, user-friendly framework for users to compare the physical associations between sRNAs and AGOs and navigate between sRNA association and functional dependence contexts without switching between disparate data sources, as illustrated in the *xol-1* and PRG-1/HRDE-1 examples described above. An future application that harnesses these advantages of AE would be to compare the extent of specificity and redundancy across the 12 WAGO-clade AGOs in *C. elegans,* which is likely to provide insights into the function of expanded AGO families in worms (Reed et al. 2020). Finally, AE’s Search and Browse modes serve complementary needs with the former supporting investigation into a specific sRNA or target of interest and the latter functioning as a hypothesis-generating tool, enabling comparative queries such as the ALG-1/ALG-2 intersection and difference operations that would otherwise require substantial bioinformatic effort.

Limitations of the current AE framework should inform how its outputs are interpreted. The enrichment and depletion thresholds applied throughout AE (minimum 5 RPM, 2-fold change, padj < 0.05) were selected to balance sensitivity and specificity but remain conservative cutoffs. Associations near these boundaries are not excluded from the underlying data but do require users to consult the “Show Evidence” plots directly. Additionally, because AE relies on IP-based enrichment rather than crosslinking-based approaches, its outputs reflect co-purification of AGO complexes with associated RNAs rather than direct, nucleotide-resolution binding evidence.

Given the broad applicability of the bioinformatic pipelines underlying AE, this database suite is poised to rapidly respond to the availability of new datasets. Incorporating additional AGO-IP or mutant depletion data requires only mapping to the relevant reference transcriptome and evaluation against the same enrichment and depletion criteria already established here. Thus, as new datasets such as wildtype *Drosophila* AGO2 with proper input control sequencing or expanded human AGO mutant backgrounds become publicly available, they can be readily incorporated into AE to close these coverage gaps.

Taken together, the Argonomics Explorer database suite transforms static, siloed AGO-sRNA datasets into a single, cross-referenced, and annotation-agnostic web platform. As demonstrated across the examples presented, AE not only recapitulates previously established AGO biology but also generates specific, testable hypotheses about AGO-cargo specificity, redundancy, and pathway crosstalk that would be challenging to uncover through manual dataset parsing.

## Materials and Methods

All high-throughput sRNA sequencing reads used to build the AE database suite were collected from the Gene Expression Omnibus (GEO) repository and Sequence Read Archive (SRA) (Clough and Barrett 2016; Seroussi et al. 2023; Fusca et al. 2025). For WAE, reads derived from Seroussi et al. (2023) and Liu et al. (2023) were aligned to a 2024 release of the *Caenorhabditis elegans* reference transcriptome (WS293) from the accredited database (Ambros et al. 2025). Reads pertaining to Argonaute-sRNA mapping were compiled from four comprehensive categories of catalogued WS293 transcripts: mRNA, non-coding RNA (ncRNA), transposons, and pseudogenic transcripts. Reads from *Drosophila melanogaster* AGO-IPs conducted by Kingston and Bartel (2021) were aligned to the reference transcriptome of R6.67, FlyBase release 2026_01, while human AGO-IP reads from Dueck et al. (2012) were mapped using release 49 (GRCh37) of the GENCODE human reference transcriptome. For both human and fly datasets, the miRbase database was used to stratify reads corresponding to mature miRNA sequences. These reference datasets were then used to map sRNA reads via Bowtie, an alignment software allowing a maximum of two mismatches (Langmead 2010). Note that since most miRNA targets only exhibit base pairing at the seed region with extensive mismatches outside this area, miRNA targets are not the focus of the AE database framework. Users interested in identifying miRNA targets can refer to other existing miRNA target databases such as mirTARCLASH and DIANA-TarBase (Vlachos et al. 2014; Yang et al. 2025).

The raw reads obtained from the corresponding datasets for each organism were preprocessed using Cutadapt to trim adapter sequences and filter the relevant transcripts based on length (16 ≤ length ≤ 30) to isolate sRNA transcripts. RPM (reads per million) normalization was then performed to remove differences in total read count between samples and enhance comparability between libraries. Unique reads were subsequently obtained by collapsing identical sequences while maintaining read counts. These processed reads were mapped to the reference transcriptome in both sense and antisense orientations. Because AGO-mediated gene regulation relies on complementary base pairing between sRNA cargo and target RNAs, the identity of functional guide strands loaded onto AGO complexes such as miRNAs corresponds to sRNA sequencing reads mapped in the sense orientation. Conversely, antisense-oriented reads reflect the target transcripts being regulated by the AGO complex.

### Immunoprecipitation-enriched reads

Datasets from two independent studies conducting systematic RIP-seq on the 19 functional *C. elegans* AGOs were analyzed and compiled in the WAE database. AGO-IP reads from WAE Dataset 1 and 2 were obtained as described in Seroussi et al. (2023) and Liu et al. (2023), respectively. Similarly, AGO-IPs and sRNA sequencing were performed in *Drosophila melanogaster* S2 cells according to Kingston and Bartel (2021) and in human HeLa cells as noted by Dueck et al. (2012).

For each AGO dataset, normalized read counts from AGO-IP samples were compared against their corresponding input controls. Enrichment was calculated as the fold change in RPM-normalized reads between IP and input samples. Significantly enriched transcripts in AGO-IP samples relative to controls were subjected to two filtering criteria before their classification as AGO-associated targets. First, the transcript must meet a minimum read count threshold of 5 RPM in IP samples to exclude low-abundance or stochastic interactions. In addition, the target transcript must exhibit at least 2-fold enrichment in the AGO-IP compared to input samples to ensure that it is preferentially pulled down with the AGO of interest. Reads that are IP-enriched in the sense orientation are considered AGO-associated sRNAs, while antisense IP-enriched reads are considered RNAs that are targeted for regulation by the corresponding AGO via sRNAs. IP-enriched small RNA data derived from this approach is consistent with previous reports on the functions of AGOs like, ALG-1, a miRNA-specific AGO in *C. elegans* (Tops et al. 2006). As expected, 91% of total ALG-1 IP-enriched reads from WAE Dataset 1 and 84% of enriched reads from WAE Dataset 2 map to miRNAs and their precursors (Figure 7). We note that targets defined by HRDE-1 and VSRA-1 IP-enriched reads derived from WAE Dataset 2 are globally inconsistent with their previously reported targets and were therefore omitted from WAE (Seroussi et al. 2023; Ding et al. 2023; Ewe et al. 2025).

**Figure 7.**
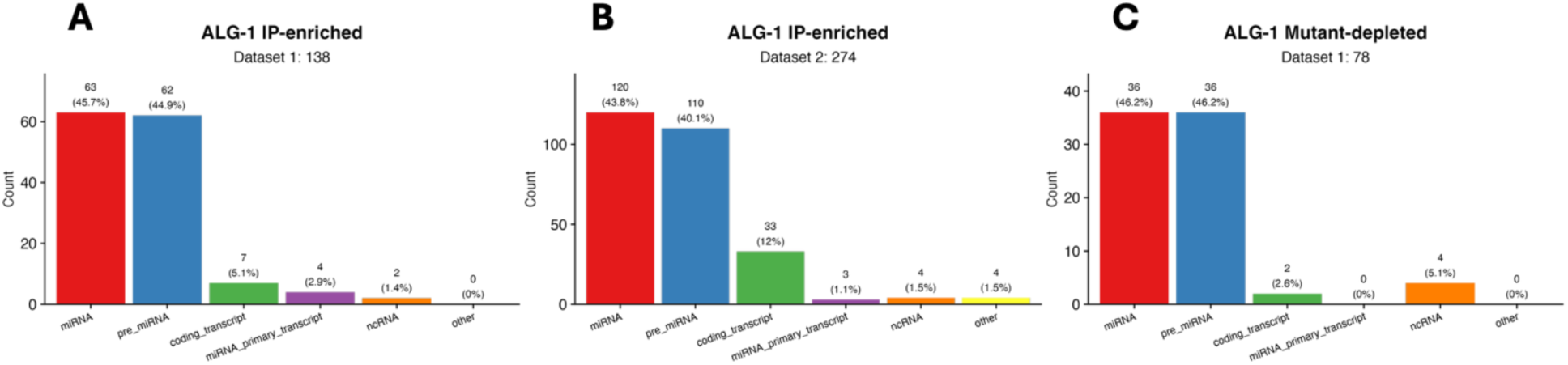
Transcript type distributions returned by WAE for the miRNA-specific AGO, ALG-1. (**7A**) Distribution of ALG-1 IP-enriched reads from Seroussi et al. 2023. (**7B**) Distribution of ALG-1 IP-enriched reads from Liu et al. 2023. (**7C**) Distribution of ALG-1 mutant-depleted reads from Seroussi et al. 2023.

### AGO mutant-depleted reads

Argonaute mutant-depleted reads for the *C. elegans* AGOs are also provided in WAE Dataset 1 and were used to identify sRNAs that rely on specific AGO proteins for their biogenesis or stability. For each AGO, sRNA profiles from wild-type samples were compared to those from corresponding AGO loss-of-function or null mutants. Depletion was quantified as significant reduction in RPM-normalized sense or antisense read abundance in AGO mutant samples relative to wildtype controls. Differential expression analysis was performed using DESeq2, and transcripts exhibiting log2 fold change below – 1 with adjusted p-values (padj) of less than 0.05 were defined as mutant-depleted targets. Mutant-depleted reads in the sense orientation correspond to the sRNAs, such as miRNAs or piRNAs, that require a specific AGO for their stability. Antisense orientation-mapped reads suggest the target RNA’s regulation depends on the mutated AGO. As expected, 92% of ALG-1 mutant-depleted reads map to miRNAs and their precursors (Figure 7). Both IP enrichment and mutant depletion datasets were included in the WAE database as these analyses provide complementary insights into AGO function. While IP enrichment captures physical association between AGOs and their targets, mutant-dependent depletion provides functional evidence that the AGO is required for maintaining the sRNA pool or target regulation.

## Acknowledgements

This work is supported in part by the National Science and Technology Council of Taiwan (NSTC 113-2221-E-006-135-MY3 and NSTC 114-2221-E-006-151-MY3) grants to W.-S.W., and the NIH grants T32GM144292 to M. B. and R35GM164060 to H.-C.L.

## Notes

https://cosbi10.ee.ncku.edu.tw/AE/

### Competing Interest Statement

The authors have declared no competing interest.

## References

1. Aalto AP, Nicastro IA, Broughton JP, Chipman LB, Schreiner WP, Chen JS, Pasquinelli AE. 2018. Opposing roles of microRNA Argonautes during Caenorhabditis elegans aging. PLoS Genet 14: e1007379. doi:10.1371/journal.pgen.1007379

2. Ambros VR, Chalfie M, Daul AL, Fire AZ, Hall DH, Horvitz HR, Mello CC, Ruvkun G, Schroeder NE, Sternberg PW, et al. 2025. From nematode to Nobel: how community-shared resources fueled the rise of *Caenorhabditis elegans* as a research organism. Proc Natl Acad Sci USA 122: e2522808122. doi:10.1073/pnas.2522808122

3. Batista PJ, Ruby JG, Claycomb JM, Chiang R, Fahlgren N, Kasschau KD, Chaves DA, Gu W, Vasale JJ, Duan S, et al. 2008. PRG-1 and 21U-RNAs interact to form the piRNA complex required for fertility in C. elegans. Mol Cell 31: 67–78. doi:10.1016/j.molcel.2008.06.002

4. Brown KC, Montgomery TA. 2018. The long and short of lifespan regulation by Argonautes. PLoS Genet 14: e1007415. doi:10.1371/journal.pgen.1007415

5. Cecere G, Hoersch S, O’Keefe S, Sachidanandam R, Grishok A. 2014. Global effects of the CSR-1 RNA interference pathway on the transcriptional landscape. Nat Struct Mol Biol 21: 358–365. doi:10.1038/nsmb.2801

6. Chakraborty S, Krishnan Y. A structural map of oncomiR-1 at single-nucleotide resolution. Nucleic Acids Research, 2017, 45(16): 9694–9705. doi:10.1093/nar/gkx613

7. Charlesworth AG, Seroussi U, Lehrbach NJ, Renaud MS, Sundby AE, Molnar RI, Lao RX, Willis AR, Woock JR, Aber MJ, et al. 2021. Two isoforms of the essential C. elegans Argonaute CSR-1 differentially regulate sperm and oocyte fertility. Nucleic Acids Res 49: 8836–8865. doi:10.1093/nar/gkab619

8. Clough E, Barrett T. 2016. The Gene Expression Omnibus Database. In: Statistical Genomics, vol 1418, 93–110. doi:10.1007/978-1-4939-3578-9_5

9. Czech B, Malone CD, Zhou R, Stark A, Schlingeheyde C, Dus M, Perrimon N, Kellis M, Wohlschlegel JA, Sachidanandam R, et al. 2008. An endogenous small interfering RNA pathway in Drosophila. Nature 453: 798–802. doi:10.1038/nature07007

10. Ding YH, Ochoa HJ, Ishidate T, Shirayama M, Mello CC. 2023. The nuclear Argonaute HRDE-1 directs target gene re-localization and shuttles to nuage to promote small RNA-mediated inherited silencing. Cell Rep 42: 112408. doi:10.1016/j.celrep.2023.112408

11. Doll MA, Soltanmohammadi N, Schumacher B. 2019. ALG-2/AGO-dependent mir-35 family regulates DNA damage-induced apoptosis through MPK-1/ERK MAPK signaling downstream of the core apoptotic machinery in Caenorhabditis elegans. Genetics 213: 173–194. doi:10.1534/genetics.119.302458

12. Dueck A, Ziegler C, Eichner A, Berezikov E, Meister G. 2012. microRNAs associated with the different human Argonaute proteins. Nucleic Acids Res 40: 9850–9862. doi:10.1093/nar/gks705

13. Ewe CK, Teichman G, Weiss S, Knott MML, Anava S, Gingold H, Sarmiento MB, Troemel E, Rechavi O. 2025. Dynamic control of Argonautes by a rapidly evolving immunological switch. Curr Biol 35: 3076–3089.e5. doi:10.1016/j.cub.2025.05.039

14. Fusca DD, Kasimatis KR, Zhu HV, Cutter AD. 2025. Dynamic birth and death of Argonaute gene family functional repertoire across Caenorhabditis nematodes. Genome Biol Evol 17: evaf106. doi:10.1093/gbe/evaf106

15. Gruszka R, Zakrzewska M. 2018. The oncogenic relevance of miR-17-92 cluster and its paralogous miR-106b-25 and miR-106a-363 clusters in brain tumors. Int J Mol Sci 19: 879. doi:10.3390/ijms19030879

16. Hutvagner G, Simard MJ. 2008. Argonaute proteins: key players in RNA silencing. Nat Rev Mol Cell Biol 9: 22–32. doi:10.1038/nrm2321

17. Jouravleva K, Zamore PD. 2025. A guide to the biogenesis and functions of endogenous small non-coding RNAs in animals. Nat Rev Mol Cell Biol 26: 347–370. doi:10.1038/s41580-024-00818-9

18. Kim VN, Han J, Siomi MC. 2009. Biogenesis of small RNAs in animals. Nat Rev Mol Cell Biol 10: 126–139. doi:10.1038/nrm2632

19. Kingston ER, Bartel DP. 2021. Ago2 protects Drosophila siRNAs and microRNAs from target-directed degradation, even in the absence of 2′-O-methylation. RNA 27: 710–724. doi:10.1261/rna.078746.121

20. Kirshner JA, Picard CL, Weiser NE, Mehta N, Feng S, Murphy VN, Vakhnovetsky A, Alessi AF, Xiao C, Inoki K, et al. 2025. Regulation of MORC-1 is key to the CSR-1–mediated germline gene licensing mechanism in C. elegans. Sci Adv 11: eado4170. doi:10.1126/sciadv.ado4170

21. Landthaler M, Gaidatzis D, Rothballer A, Chen PY, Soll SJ, Dinic L, Ojo T, Hafner M, Zavolan M, Tuschl T. 2008. Molecular characterization of human Argonaute-containing ribonucleoprotein complexes and their bound target mRNAs. RNA 14: 2580–2596. doi:10.1261/rna.1351608

22. Langmead B. 2010. Aligning short sequencing reads with Bowtie. Curr Protoc Bioinforma 32.

23. Lee HC, Gu W, Shirayama M, Youngman E, Conte D Jr, Mello CC. 2012. C. elegans piRNAs mediate the genome-wide surveillance of germline transcripts. Cell 150: 78–87. doi:10.1016/j.cell.2012.06.016

24. Lewis BP, Shih IH, Jones-Rhoades MW, Bartel DP, Burge CB. 2003. Prediction of mammalian microRNA targets. Cell 115: 787–798. doi:10.1016/S0092-8674(03)01018-3

25. Liu L, Wang X, Zhao W, Li Q, Li J, Chen H, Shan G. 2023. Systematic characterization of small RNAs associated with C. elegans Argonautes. Sci China Life Sci 66: 1303–1322. doi:10.1007/s11427-022-2304-8

26. Meyer WJ, Schreiber S, Guo Y, Volkmann T, Welte MA, Müller HAJ. 2006. Overlapping functions of Argonaute proteins in patterning and morphogenesis of Drosophila embryos. PLoS Genet 2: e134. doi:10.1371/journal.pgen.0020134

27. Nakanishi K. 2022. Anatomy of four human Argonaute proteins. Nucleic Acids Res 50: 6618–6638. doi:10.1093/nar/gkac519

28. Okamura K, Liu N, Lai EC. 2009. Distinct mechanisms for microRNA strand selection by Drosophila Argonautes. Mol Cell 36: 431–444. doi:10.1016/j.molcel.2009.09.027

29. Pal A, Vasudevan V, Houle F, Lantin M, Maniates KA, Quévillon Huberdeau M, Abbott AL, Simard MJ. 2024. Defining the contribution of microRNA-specific Argonautes with slicer capability in animals. Nucleic Acids Res 52: 5002–5015. doi:10.1093/nar/gkae173

30. Reed KJ, Svendsen JM, Brown KC, Montgomery BE, Marks TN, Vijayasarathy T, Parker DM, Osborne Nishimura E, Updike DL, Montgomery TA. 2020. Widespread roles for piRNAs and WAGO-class siRNAs in shaping the germline transcriptome of Caenorhabditis elegans. Nucleic Acids Res 48: 1811–1827. doi:10.1093/nar/gkz1178

31. Seroussi U, Lugowski A, Wadi L, Lao RX, Willis AR, Zhao W, Sundby AE, Charlesworth AG, Reinke AW, Claycomb JM. 2023. A comprehensive survey of C. elegans argonaute proteins reveals organism-wide gene regulatory networks and functions. eLife 12: e83853. doi:10.7554/eLife.83853

32. Shen EZ, Chen H, Ozturk AR, Tu S, Shirayama M, Tang W, Ding YH, Dai SY, Weng Z, Mello CC. 2018. Identification of piRNA binding sites reveals the Argonaute regulatory landscape of the C. elegans germline. Cell 172: 937–951.e18. doi:10.1016/j.cell.2018.02.002

33. Shirayama M, Seth M, Lee HC, Gu W, Ishidate T, Conte D Jr, Mello CC. 2012. piRNAs initiate an epigenetic memory of nonself RNA in the C. elegans germline. Cell 150: 65–77. doi:10.1016/j.cell.2012.06.015

34. Tang W, Seth M, Tu S, Shen EZ, Li Q, Shirayama M, Weng Z, Mello CC. 2018. A sex chromosome piRNA promotes robust dosage compensation and sex determination in C. elegans. Dev Cell 44: 762–770.e3. doi:10.1016/j.devcel.2018.01.025

35. Tops BBJ, Plasterk RHA, Ketting RF. 2006. The Caenorhabditis elegans Argonautes ALG-1 and ALG-2: almost identical yet different. Cold Spring Harb Symp Quant Biol 71: 189–194. doi:10.1101/sqb.2006.71.035

36. Vlachos IS, Paraskevopoulou MD, Karagkouni D, Georgakilas G, Vergoulis T, Kanellos I, Anastasopoulos IL, Maniou S, Karathanou K, Kalfakakou D, et al. 2015. DIANA-TarBase v7.0: indexing more than half a million experimentally supported miRNA:mRNA interactions. Nucleic Acids Res 43: D153–D159. doi:10.1093/nar/gku1215

37. Yang TH, Li XW, Lee YH, Lu SY, Wu WS, Lee HC. 2025. mirTarCLASH: a comprehensive miRNA target database based on chimeric read-based experiments. Database 2025: baaf023. doi:10.1093/database/baaf023

38. Yigit E, Batista PJ, Bei Y, Pang KM, Chen CC, Tolia NH, Joshua-Tor L, Mitani S, Simard MJ, Mello CC. 2006. Analysis of the C. elegans Argonaute family reveals that distinct Argonautes act sequentially during RNAi. Cell 127: 747–757. doi:10.1016/j.cell.2006.09.033

