## Supplementary Figures for "Argonomics Explorer: an interactive database for dissecting Argonaute-RNA regulatory networks"

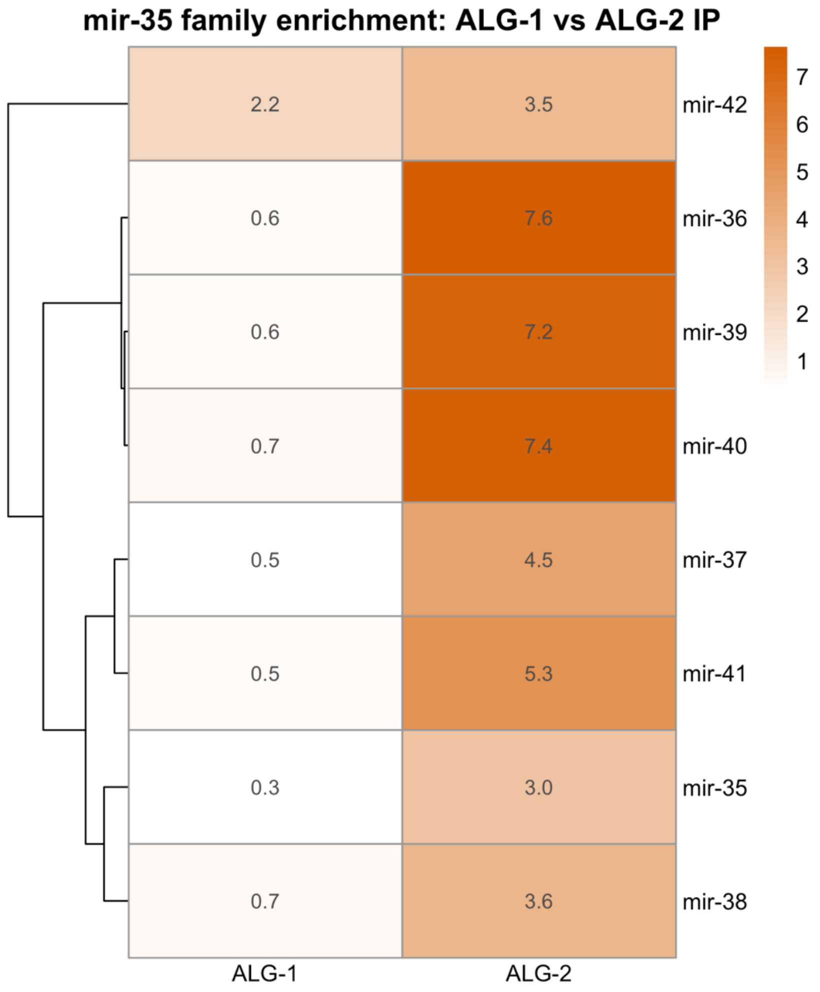


**Supplementary Figure S1**

*mir-35* family member enrichment in ALG-1 and ALG-2 IP. Heatmap of $log_2$fold-change values (average of two replicates, WAE Dataset 1; Seroussi et al. 2023) for each *mir-35-42* family member in ALG-1 an ALG-2 IP. Rows are hierarchically clustered on column-scaled (z-scored) enrichment values so that ALG-1 and ALG-2 contribute equally to clustering distance. Displayed values and colors reflect raw, unscaled enrichment. This clustering separates *mir-42,* which shows comparable enrichment in both AGOs, from the other seven family members, which cluster together with strong ALG-2 association preference.

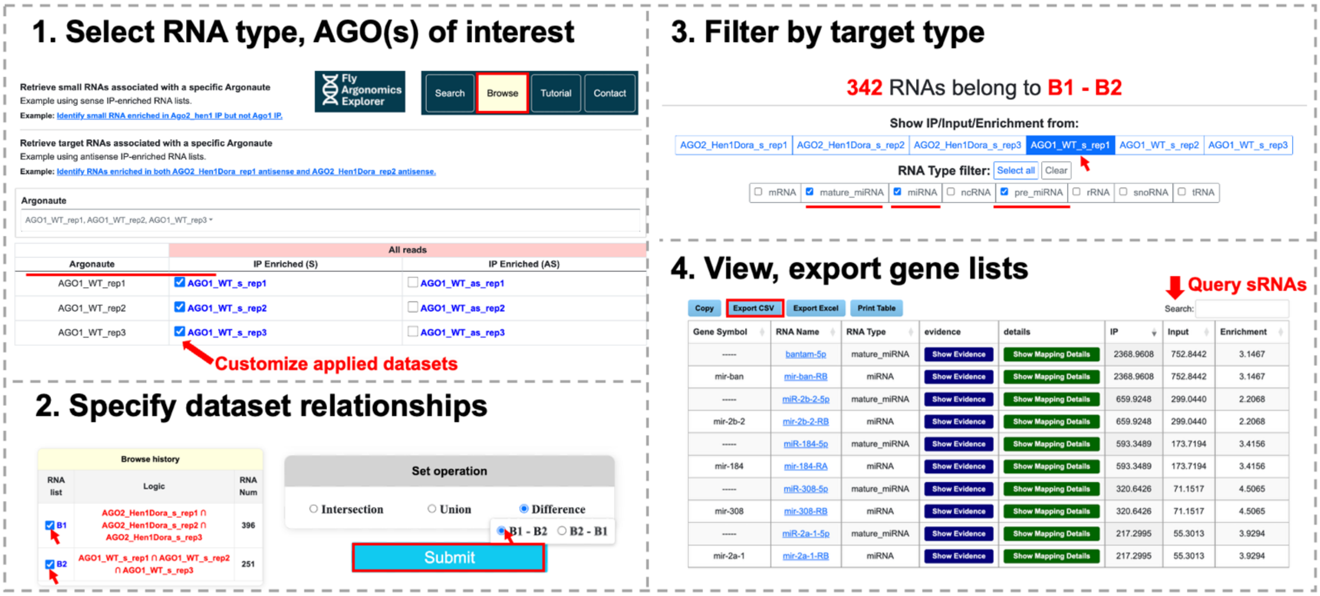


### Supplementary Figure S2

Generating miRNA lists that preferentially associated with AGO-2 but not AGO-1 in Fly Argonomics Explorer (FAE) Browse mode. Specify AGO(s) of interest and reads used for mapping (**S2.1**)**.** Apply set operations to selected datasets, retaining access to past queries (**S2.2**). Select sRNA categories of interest and choose display dataset for the resulting table (**S2.3**). View and export sRNA tables containing gene name, RNA type, and IP, input, and enrichment values from the specified dataset (**S2.4**). Note that bantam* and miR-184 are among selected miRNA* strands are specifically sorted to AGO2.

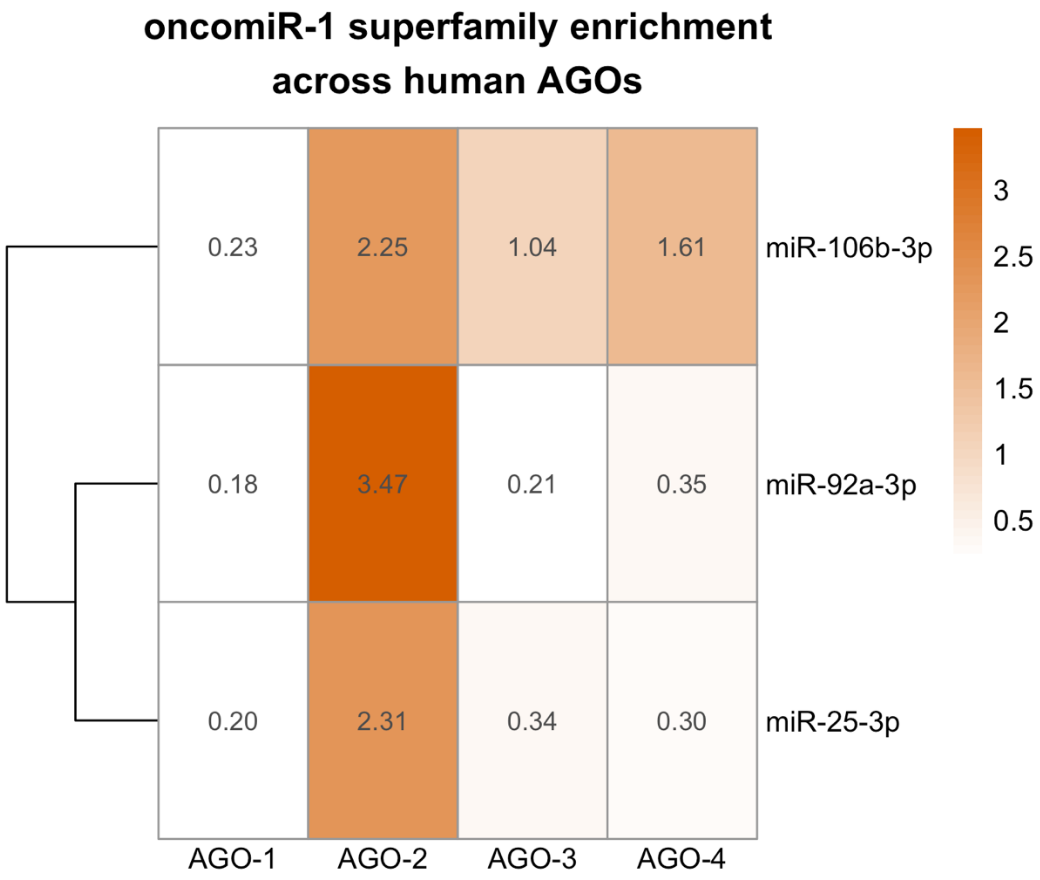


**Supplementary Figure S3**

oncomiR-1 mature miRNA enrichment across human AGOs. Heatmap of enrichment values (IP relative to input) for one representative mature miRNA from each of the three oncomiR-1 paralogous clusters – miR-92a-3p (miR-17-92), miR-25-3p (miR-106a-363), and miR-106b-3p (miR-106b-25) – across AGO-1, AGO-2, AGO-3, and AGO-4 IP. Rows are hierarchically clustered on column-scaled enrichment values; displayed values and colors reflect raw, unscaled enrichment. All three paralogs show preferential enrichment in AGO-2 relative to the other three AGOs, consistent with coordinated AGO-2 loading across the oncomiR-1 superfamily.
